# AVDB: The Arab Variation and Disease Burden Database

**DOI:** 10.1101/2025.08.24.671996

**Authors:** Omer S. Alkhnbashi, Ruchi Jain, Sathishkumar Ramaswamy, Costerwell Khyriem, Sven Hauns, Rasheed Mohammad, Rolf Backofen, Ahmad Abou Tayoun

## Abstract

**Motivation:** Public genomic databases are crucial to precision medicine but often lack representation from Arab populations, which have distinct genetic structures due to high consanguinity rates.

**Results:** We present the Arab Variation and Disease Burden Database (AVDB), derived from 1,194 exomes of Emirati individuals, comprising 2,481 curated variants across 850 genes. AVDB provides pathogenicity classification, carrier frequencies, and gene-level at-risk couple rates (ARCR), reaching 21% under first-cousin mating models. It integrates machine learning for gene prioritization and offers a customizable panel builder. Compared to global panels, AVDB captures more regional burden, filling a major gap in population-specific screening. The resource supports genomic diagnostics and health policy for Arab populations and is freely accessible at https://avdb-arabgenome.ae.

**Supplementary information:** Supplementary data are available at *Bioinformatics* online.

## 1 Introduction

A comprehensive and representative genomic database is essential for accurate variant interpretation, disease gene discovery, and the development of effective population screening programs [1]. However, existing public resources such as gnomAD [2], ClinVar [3], and ClinGen [4] show a significant bias towards populations of European ancestry, with limited representation of Middle Eastern cohorts [5]. This lack of diversity reduces the usefulness of these databases for underrepresented groups, especially in regions like the Arabian Peninsula, where the genetic landscape is shaped by distinct historical, cultural, and demographic factors [6].

Arab populations, particularly those in the Gulf region, exhibit some of the highest global rates of consanguinity, leading to a considerable burden of autosomal recessive disorders [7]. Although initiatives like the Greater Middle East (GME) Variome Project [8] and the Iranome Project [9] have made valuable strides in cataloguing regional variation, these efforts remain either inaccessible to clinicians or limited in population coverage, phenotype integration, or clinical annotations. Consequently, there remains a pressing need for publicly accessible, clinically curated databases tailored to Arab genomic data to support both research and clinical utility. To address this gap, we have developed the Arab Variation and Disease [5,6] Burden Database (AVDB), a curated and open-access genomic resource based on exome data from 1,194 unrelated Emirati individuals [1]. AVDB extends beyond frequency-based variant repositories by incorporating expert-curated annotations, variant-level pathogenicity classification, gene-level carrier frequencies, population-specific at-risk couple rates (ARCR), and machine learning–based gene prioritization. The platform also includes a customizable screening panel builder designed for clinical and public health applications.

Unlike previous regional efforts, AVDB is the first fully open-access, Arab-specific database that links clinical variant interpretation with disease burden metrics. Hosted at the Dubai Health Genomic Medicine Centre, AVDB serves as a scalable framework for precision screening in consanguineous populations and offers an essential reference for researchers, clinicians, and policymakers working toward equitable genomic medicine [1].

## 2 Methods

Between 2020 and 2024, a total of 1,194 unrelated Emirati individuals underwent clinical exome sequencing [1]. The DNA samples were fragmented using ultrasonication (Covaris), then captured with Agilent Clinical Research Exome v2 probes, and sequenced on the Illumina NovaSeq 6000 platform (2×150 bp, with an average depth of ≥100X). The sequencing reads were aligned to the GRCh37 reference genome using BWA [10], and variants were jointly called with bcftools version 1.19 [11]. The annotations at the variant level included RefSeq transcripts [12], gnomAD allele frequencies [2], and ClinVar pathogenicity assertions [3]. The AVDB data processing workflow, including sequencing, annotation, filtering, variant classification, and database outputs, is summarized in Figure 1.

**Figure 1.**
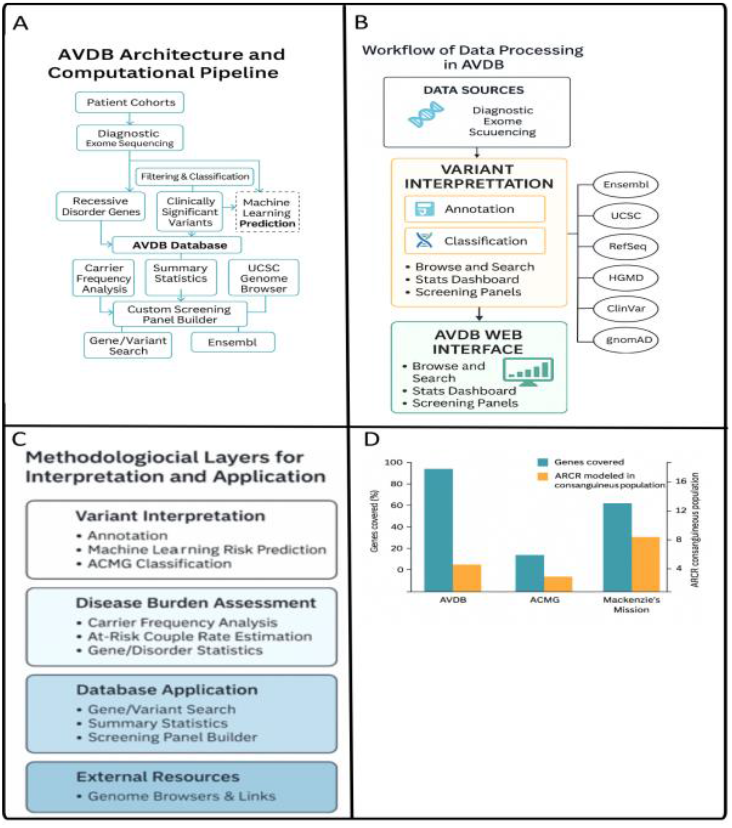
Overview of the AVDB platform and data processing pipeline. **(A)** *AVDB system architecture and Computational Pipeline*: A schematic representation showing the integration of diagnostic exome sequencing data from patient cohorts into the AVDB backend. The platform supports variant filtering, ACMG-guided classification, curation, and storage in a structured database accessible through a user-friendly web interface. **(B)** *Workflow of data processing in AVDB*: The core pipeline includes variant calling, annotation, and pathogenicity classification, followed by integration into population-specific modules. Annotation sources include *Ensembl, UCSC Genome Browser, gnomAD, RefSeq, ClinVar*, and *HGMD*. Data outputs support variant- and gene-level statistics, carrier rates, and gene panel customization. **(C)** *Methodological framework*: AVDB’s design incorporates expert variant curation, phenotype-linked interpretation, and machine learning-based gene scoring to support downstream clinical decision-making and population screening strategies (D) *Comparative interpretation of AVDB-derived gene burden*, showing how AVDB captures disease-relevant gene content unmatched by international reproductive screening panels. The 113-gene ACMG panel detects only 56 of the 425 AVDB genes, modeling an ARCR of just 6%. Mackenzie’s Mission (∼1,300 genes) improves coverage to 76% of AVDB genes with a modeled ARCR of 12.4%. However, 24% of clinically relevant AVDB genes remain absent, highlighting that global panels may inadequately represent populations with distinct genetic architectures such as the Emiratis.

### 2.1 Variant Filtering and Pathogenicity Assignment

To ensure accuracy and clinical relevance, a multi-step variant filtering and classification protocol was implemented. Initially, variants were limited to high-confidence, protein-coding changes with adequate sequencing coverage (see Figure 1A). Only variants with a minimum read depth of 10× and a Phred quality score exceeding 30 were retained for further analysis. Additionally, the focus was narrowed to exonic regions and intronic boundaries extending ±8 base pairs from exon junctions, in accordance with RefSeq transcript annotations [12].

Following the technical filtering phase, pathogenicity was assessed using a hierarchical framework based on guidelines established by the American College of Medical Genetics and Genomics and the Association for Molecular Pathology (ACMG/AMP) [13]. ClinVar [3] served as a primary reference database, accepting variants with two or more submitters and a review status of at least two stars, classified as either “Pathogenic” or “Likely Pathogenic” without reclassification. For variants not conclusively characterized in ClinVar, in-house clinical variant scientists conducted comprehensive ACMG/AMP interpretation utilizing VarSome [14] and manual literature review, incorporating functional predictions and population frequency thresholds.

Furthermore, rare novel protein-truncating variants, such as stop-gain, frameshift, and canonical splice-site mutations, were deemed Likely Pathogenic if they satisfied the PVS1 and other evidence criteria according to the ACMG/AMP guideline [15]. Only genes confirmed to exhibit autosomal recessive inheritance were included, and gene-disease associations were cross-verified utilizing ClinGen [4], OMIM [16], the ACMG Secondary Findings v3.0 list [17], and Mackenzie’s Mission screening panel [18] to ensure clinical validity and actionability.

This comprehensive approach facilitated the creation of a curated set of pathogenic and likely pathogenic variants that accurately represent the Emirati population, rendering the dataset suitable for subsequent carrier frequency estimation and screening burden analysis.

### Platform Infrastructure, Accessibility, and Data Availability

AVDB is described as a modular database platform that combines robust technical architecture with an intuitive user interface aimed at population-scale variant interpretation and reproductive risk modelling. The backend is built using Python with the Django framework and connects to a PostgreSQL relational database that stores curated variants, gene-level statistics, and ARCR models (see Figure 1B). The frontend, powered by React.js, offers a responsive, browser-independent interface that enables smooth user interactions. The entire system is hosted on secure cloud infrastructure managed by the Dubai Health Genomic Medicine Centre, ensuring high availability and scalability. The platform seamlessly integrates curated data with external resources, including gnomAD [2], ClinVar [3], RefSeq [12], UCSC genome browser [19] and Ensembl [20], through automated pipelines that facilitate scheduled quarterly data updates. All annotations and scoring are systematically organized within structured schemas designed for high-speed querying across gene, variant, and disease categories.

AVDB has undergone comprehensive testing across modern web browsers, including Chrome, Firefox, Safari, and Edge, and demonstrates full responsiveness on both desktop and tablet devices. Users can access all features of the site without needing to register. The platform offers multiple data export options, allowing users to download gene- and variant-level statistics such as CSV files directly from their browsers. Additionally, a RESTful API is planned for implementation in upcoming updates to improve integration with diagnostic pipelines and third-party applications.

All data, including curated variant lists and population-specific metrics, will remain publicly accessible for at least two years following publication. Supplementary materials such as screenshots, extended gene tables, and user guides that are available on the website and cited in Supplementary Data 1.

## 3 Results

### 3.1 Variant Landscape

In a detailed analysis of 1,194 unrelated Emirati exomes, we identified a total of 600,179 protein-coding and near-splice variants. Among these, 799 unique variants were classified as pathogenic or likely pathogenic (P/LP) based on the guidelines established by the American College of Medical Genetics and Genomics (ACMG) and the Association for Molecular Pathology (AMP). These variants were spread across 425 genes linked to autosomal recessive diseases, which formed the main focus for subsequent analyses. Interestingly, 130,431 variants, making up 21.7% of the entire dataset, were absent from the latest release of the Genome Aggregation Database (gnomAD), emphasizing the significant level of population-specific variation within the Emirati cohort. The structure of the AVDB database and its core functionalities, including gene- and variant-level outputs, are depicted in Figure 1.

### 3.2 Features and Functionality

AVDB offers a comprehensive suite of interactive modules designed to support clinicians, researchers, and public health stakeholders in analyzing population-specific variations, estimating carrier burdens, and personalizing screening strategies. Each module within the platform is specifically designed to enhance usability and clinical relevance, reflecting the unique genetic traits of Arab populations (See Figure 1D).

The platform is mainly driven by an advanced search engine that supports queries using gene symbols (HGNC), transcript IDs, variant HGVS nomenclature, or disease-related keywords. Users receive a structured output that includes curated pathogenic and likely pathogenic (P/LP) variants, as well as ACMG classifications. The platform also offers links to trusted external databases like ClinVar, gnomAD, and Ensembl. Variant-level frequencies specific to the Emirati population are highlighted, along with gene-level summaries that include cumulative carrier frequency and identified founder mutations.

### 3.3 Screening Burden and Panel Comparison

Utilizing the curated gene-level data, we assessed the cumulative at-risk couple rate (ARCR) under various population mating models. In scenarios involving random mating among Emiratis, the ARCR for the 425 identified genes was estimated to be approximately 4%. However, under a first cousin mating scenario prevalent cultural practice in the region; the ARCR increased significantly to 21%. These estimates suggest that one in five consanguineous Emirati couples may theoretically be at increased risk of conceiving a child affected by an autosomal recessive condition detectable through the AVDB (See Figure 1D).

To contextualize these findings, we compared the carrier burden identified through the AVDB with several established reproductive screening panels. The 113-gene ACMG panel, which is widely recommended for expanded carrier screening, captured only 56 of the 425 AVDB genes and yielded a notably lower ARCR of 6% in the context of consanguineous mating. The Mackenzie’s Mission panel, encompassing over 1,300 genes, demonstrated improved performance, covering approximately 76% of the AVDB genes though producing a reduced ARCR of 12.4% relative to AVDB under first-cousin mating assumption. This finding indicates that even the most comprehensive international screening panels may be insufficient for populations with unique genetic architectures, such as the Emiratis (See Figure 1D).

Furthermore, when comparing the AVDB gene list to the 570-gene panel currently recommended by the UAE Ministry of Health and Prevention (MOHAP), it was observed that only 50% of the top 100 high-burden genes were included. This disparity emphasizes the limitations of establishing regional screening panels based on global variant databases. It underlines the importance of AVDB as a population-specific evidence base.

## 4 Discussion and Conclusion

Arab populations face a unique challenge: while high rates of consanguinity significantly increase the prevalence of recessive disorders, most global genomic databases offer limited insight into region-specific pathogenic variants. Although prior efforts such as the Greater Middle East Variome Project and Iranome have catalogued regional variation, these initiatives often lack open-access clinical annotations, integrated disease burden models, or direct applicability for public health screening programs.

The Arab Variation and Disease Burden Database (AVDB) addresses these gaps by offering a clinically curated, population-specific, and openly accessible resource tailored for Emirati and Arab populations. In contrast to the gnomAD or ClinVar repositories, which focus primarily on variant frequency or aggregated submissions, AVDB provides curated pathogenicity assessments using ACMG/AMP criteria, gene-level carrier frequency estimates, and modeled at-risk couple rates under culturally relevant mating patterns such as first-cousin unions. Compared to widely adopted panels such as the ACMG 113-gene panel or Mackenzie’s Mission, AVDB captures a broader spectrum of pathogenic variation observed in the Emirati population, identifying nearly 50% more high-burden genes not represented in those panels.

From a clinical perspective, AVDB provides variant-level insights and gene prioritization that can be directly integrated into diagnostic pipelines for rare disease patients. For public health stakeholders and policymakers, the platform offers actionable metrics such as carrier burden estimates and customizable screening panels, enabling evidence-based updates to national screening programs such as the UAE’s premarital testing strategy. For example, AVDB highlights key high-burden genes like *DONSON* and *ALDH3A2* that are currently absent from national screening guidelines but present significant risk within the local population.

Despite its contributions, AVDB has certain limitations. The current dataset is limited to exome sequencing from 1,194 Emirati individuals and may not fully capture variation in non-coding regions or large structural variants. As such, future versions of AVDB will integrate whole-genome sequencing (WGS) data and long-read technologies to improve detection of complex genomic features such as *CYP21A2* pseudogene rearrangements. In addition, the platform will expand to include data from other Arab populations to enhance regional representation and support cross-country genomic comparisons. Incorporating functional genomics data and phenotype-genotype correlations from local clinics will further strengthen the clinical utility of AVDB.

In conclusion, AVDB represents a scalable and clinically actionable model for genomic resource development in underrepresented populations. By offering a high-resolution, curated, and locally relevant database, it not only improves variant interpretation in clinical diagnostics but also informs population screening strategies and public health decision-making across the Arab world.

## Supporting information

https://avdb-arabgenome.ae

## Acknowledgements

The authors sincerely thank the families and patients whose anonymized genomic data supported this research. We also appreciate the clinical geneticists, laboratory scientists, and bioinformaticians at Genomic Medicine Centre for their vital contributions to variant curation and platform assessment. Additionally, we recognize the efforts of the AVDB development team in frontend design, database integration, and deployment, which were essential to this project’s success.

## References

[1] Jain R, Bizzari S, Ramaswamy S, et al. Pathogenic variation underlying rare diseases in an Arab population: implications for screening programs. Genetics in Medicine Open. 2025; doi:10.1016/j.gimo.2025.103446.

[2] Karczewski KJ, Francioli LC, Tiao G, et al. The mutational constraint spectrum quantified from variation in 141,456 humans. Nature. 2020;581:434–443. doi:10.1038/s41586-020-2308-7.

[3] Landrum MJ, Lee JM, Riley GR, et al. ClinVar: public archive of relationships among sequence variation and human phenotype. Nucleic Acids Res. 2014;42:D980–D985. doi:10.1093/nar/gkt1113.

[4] Rehm HL, Berg JS, Brooks LD, et al. ClinGen—the Clinical Genome Resource. N Engl doi:10.1056/NEJMsr1406261. J Med. 2015;372:2235–2242.

[5] Scott EM, Halees A, Itan Y, et al. Characterization of Greater Middle Eastern genetic variation for enhanced disease gene discovery. Nat Genet. 2016;48(9):1071–1076. doi:10.1038/ng.3592.

[6] Hajjej A, Almawi WY, Arnaiz-Villena A, et al. HLA genes in the Arab world: a narrative review. PLoS One. 2018;13(3):e0192269. doi:10.1371/journal.pone.0192269.

[7] Al-Gazali L, Hamamy H, Al-Arrayad S. Genetic disorders in the Arab world. BMJ. 2006;333:831–834. doi:10.1136/bmj.38982.704931.AE.

[8] Scott EM, et al. (same as [5]; GME Variome).

[9] Fattahi Z, Beheshtian M, Mohseni M, et al. Iranome: a catalog of genomic variations in the Iranian population. Hum Mutat. 2019;40(11):1968–1984. doi:10.1002/humu.23880.

[10] Li H, Durbin R. Bioinformatics. 2009;25(14):1754–1760. doi:10.1093/bioinformatics/btp324.

[11] Danecek P, Bonfield JK, Liddle J, et al. Gigascience. 2021;10(2):giab008. doi:10.1093/gigascience/giab008.

[12] O’Leary NA, Wright MW, Brister JR, et al. Nucleic Acids Res. 2016;44(D1):D733–D745. doi:10.1093/nar/gkv1189.

[13] Richards S, Aziz N, Bale S, et al. Genet Med. 2015;17:405–424. doi:10.1038/gim.2015.30.

[14] Kopanos C, Tsiolkas V, Kouris A, et al. Bioinformatics. 2019;35(11):1974–1976. doi:10.1093/bioinformatics/bty897.

[15] Abou Tayoun AN, Pesaran T, DiStefano MT, et al. Hum Mutat. 2018;39(11):1517–1524. doi:10.1002/humu.23626.

[16] Amberger JS, Bocchini CA, Scott AF, Hamosh A. Nucleic Acids Res. 2019;47(D1):D1038–D1043. doi:10.1093/nar/gky1151.

[17] Miller DT, Lee K, Gordon AS, et al. Genet Med. 2021;23:1391– 1398. doi:10.1038/s41436-021-01171-4.

[18] Kirk EP, Barlow-Stewart K, Selvanathan A, et al. Genet Med. 2022;24:733–743. doi:10.1016/j.gim.2021.12.004.

[19] Kent WJ, Sugnet CW, Furey TS, et al. Genome Res. 2002;12(6):996–1006. doi:10.1101/gr.229102.

[20] Yates AD, Allen J, Amode RM, et al. Nucleic Acids Res. 2020;48(D1):D682–D688. doi:10.1093/nar/gkz966.

[21] Abou Tayoun AN. Nat Rev Genet. 2023;24(12):801–802. doi:10.1038/s41576-023-00654-1.

[22] Tayoun AA, Sinha S, Rabea F, et al. Research Square (preprint). 2024. doi:10.21203/rs.3.rs-4235049/v1.

[23] Hall B, Alyafei S, Ramaswamy S, et al. medRxiv. 2024. doi:10.1101/2024.02.22.24303180.

