## Supplementary material for "AVDB: The Arab Variation and Disease Burden Database": https://avdb-arabgenome.ae

Supplementary Data 1 – Arab Variation and Disease Burden Database (AVDB)

1. Platform Screenshots

   The following screenshots illustrate the core functionalities of the AVDB web platform (https://avdb-arabgenome.ae), highlighting how users can search, visualize, and export population-specific genomic data.

   - Home page and search bar

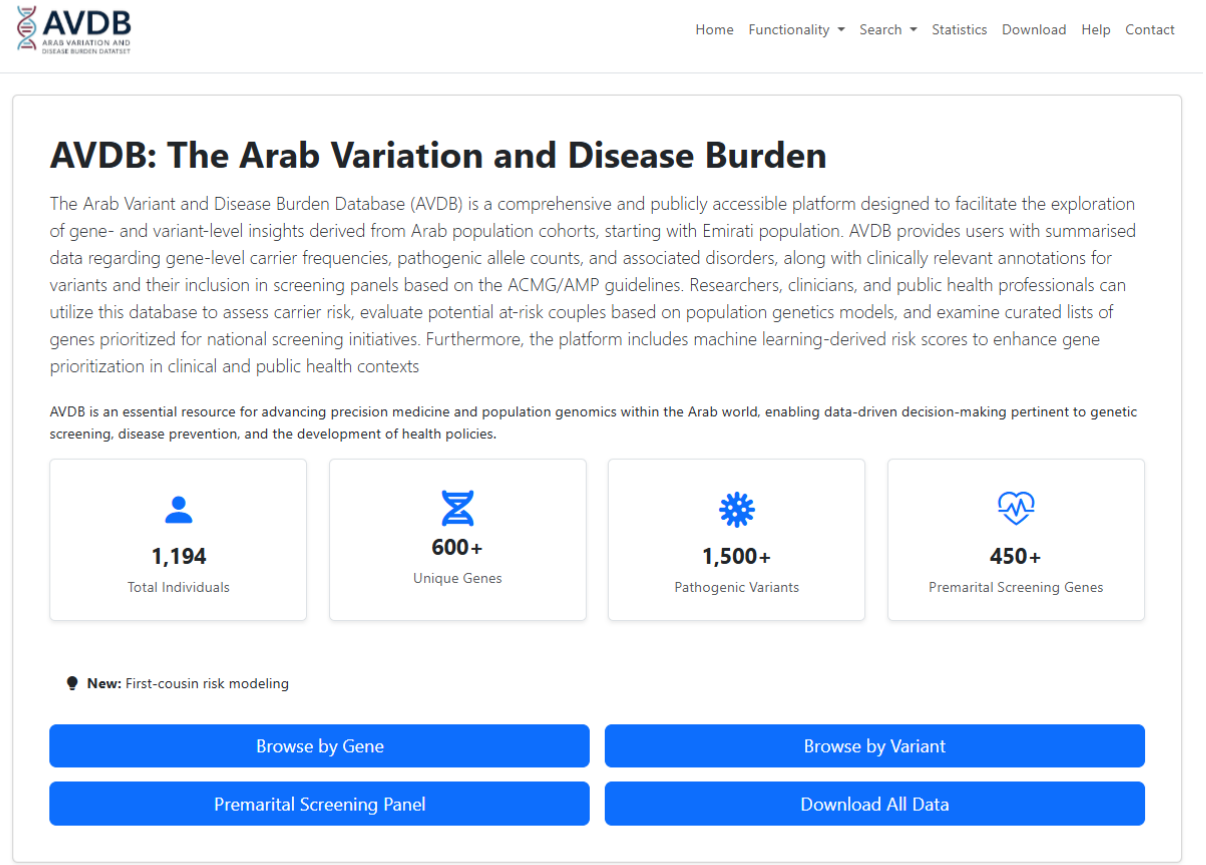

AVDB ARAB VARIATION AND DISEASE BURDEN DATASET

Home Functionality Search Statistics Download Help Contact

### AVDB: The Arab Variation and Disease Burden

The Arab Variant and Disease Burden Database (AVDB) is a comprehensive and publicly accessible platform designed to facilitate the exploration of gene- and variant-level insights derived from Arab population cohorts, starting with Emirati population. AVDB provides users with summarised data regarding gene-level carrier frequencies, pathogenic allele counts, and associated disorders, along with clinically relevant annotations for variants and their inclusion in screening panels based on the ACMG/AMP guidelines. Researchers, clinicians, and public health professionals can utilize this database to assess carrier risk, evaluate potential at-risk couples based on population genetics models, and examine curated lists of genes prioritized for national screening initiatives. Furthermore, the platform includes machine learning-derived risk scores to enhance gene prioritization in clinical and public health contexts

AVDB is an essential resource for advancing precision medicine and population genomics within the Arab world, enabling data-driven decision-making pertinent to genetic screening, disease prevention, and the development of health policies.

1,194 Total Individuals

600+ Unique Genes

1,500+ Pathogenic Variants

450+ Premarital Screening Genes

**New:** First-cousin risk modeling

Browse by Gene | Browse by Variant

Premarital Screening Panel | Download All Data

   - Gene search results

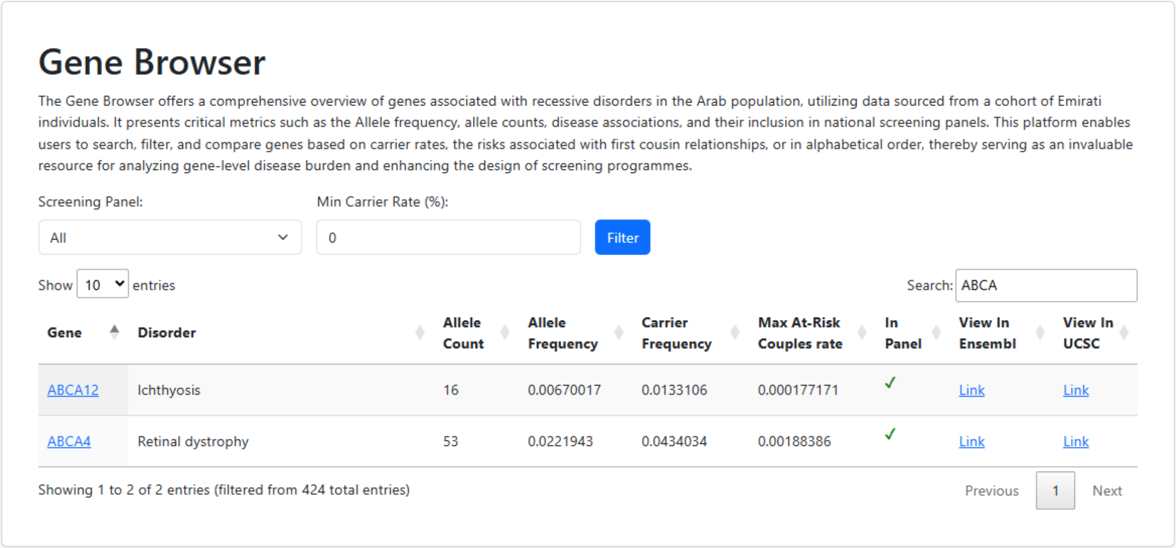

### Gene Browser

The Gene Browser offers a comprehensive overview of genes associated with recessive disorders in the Arab population, utilizing data sourced from a cohort of Emirati individuals. It presents critical metrics such as the Allele frequency, allele counts, disease associations, and their inclusion in national screening panels. This platform enables users to search, filter, and compare genes based on carrier rates, the risks associated with first cousin relationships, or in alphabetical order, thereby serving as an invaluable resource for analyzing gene-level disease burden and enhancing the design of screening programmes.

Screening Panel: All

Min Carrier Rate (%): 0

Filter

Show 10 entries

Search: ABCA

| Gene | Disorder | Allele Count | Allele Frequency | Carrier Frequency | Max At-Risk Couples rate | In Panel | View In Ensembl | View In UCSC |
|---|---|---|---|---|---|---|---|---|
| ABCA12 | Ichthyosis | 16 | 0.00670017 | 0.0133106 | 0.000177171 | ✓ | Link | Link |
| ABCA4 | Retinal dystrophy | 53 | 0.0221943 | 0.0434034 | 0.00188386 | ✓ | Link | Link |

Showing 1 to 2 of 2 entries (filtered from 424 total entries)

Previous 1 Next

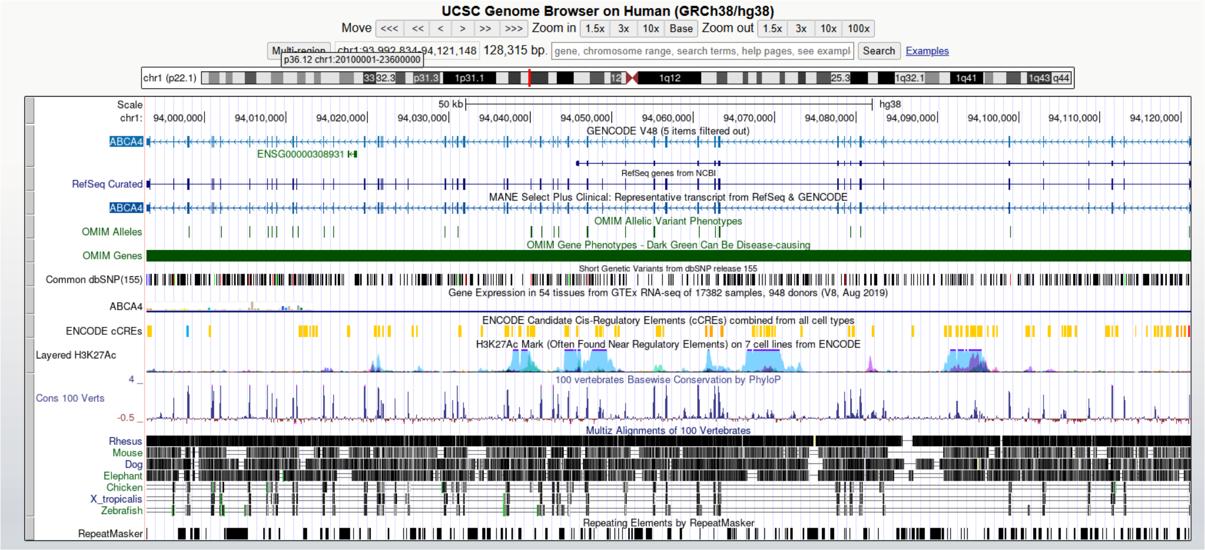
UCSC Genome Browser on Human (GRCh38/hg38)
Move <<< << < > >> >>> Zoom in 1.5x 3x 10x Base Zoom out 1.5x 3x 10x 100x
Multi-region chr1:93,992,834-94,121,148 128,315 bp. gene, chromosome range, search terms, help pages, see exampl Search Examples
p36.12 chr1:20100001-23600000
chr1 (p22.1)
Scale
chr1:
50 kb
hg38
94,000,000 94,010,000 94,020,000 94,030,000 94,040,000 94,050,000 94,060,000 94,070,000 94,080,000 94,090,000 94,100,000 94,110,000 94,120,000
GENCODE V48 (5 items filtered out)
ABCA4
ENSG00000308931
RefSeq genes from NCBI
RefSeq Curated
MANE Select Plus Clinical: Representative transcript from RefSeq & GENCODE
ABCA4
OMIM Allelic Variant Phenotypes
OMIM Alleles
OMIM Gene Phenotypes - Dark Green Can Be Disease-causing
OMIM Genes
Short Genetic Variants from dbSNP release 155
Common dbSNP(155)
Gene Expression in 54 tissues from GTEx RNA-seq of 17382 samples, 948 donors (V8, Aug 2019)
ABCA4
ENCODE Candidate Cis-Regulatory Elements (cCREs) combined from all cell types
ENCODE cCREs
H3K27Ac Mark (Often Found Near Regulatory Elements) on 7 cell lines from ENCODE
Layered H3K27Ac
100 vertebrates Basewise Conservation by PhyloP
4
Cons 100 Verts
-0.5
Multiz Alignments of 100 Vertebrates
Rhesus
Mouse
Dog
Elephant
Chicken
X_tropicalis
Zebrafish
Repeating Elements by RepeatMasker
RepeatMasker

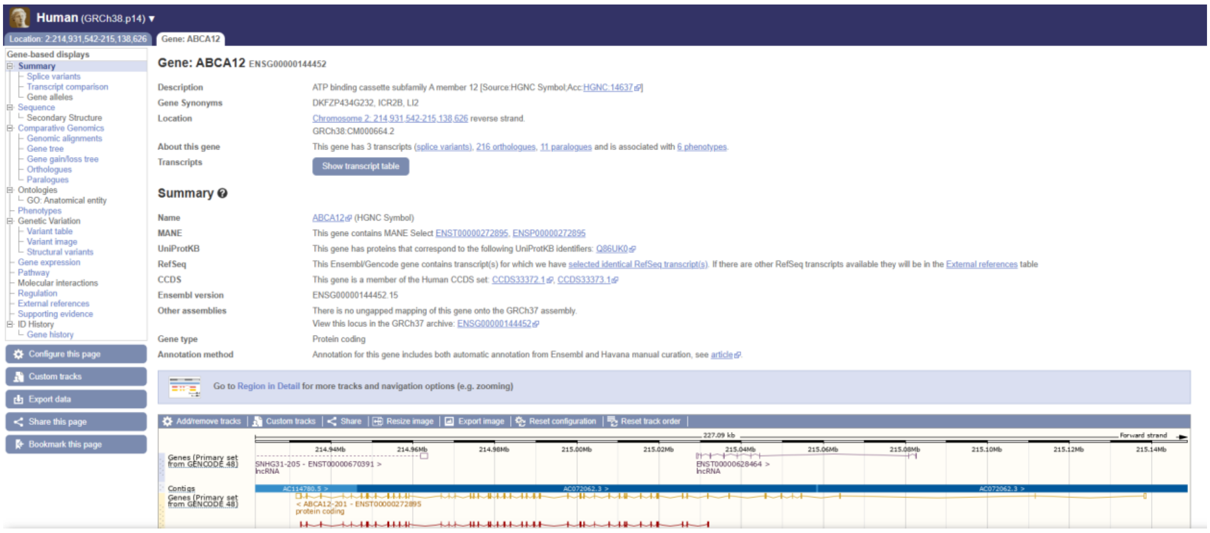
Human (GRCh38.p14)
Location: 2:214,931,542-215,138,626
Gene: ABCA12
Gene-based displays
Summary
Splice variants
Transcript comparison
Gene alleles
Sequence
Secondary Structure
Comparative Genomics
Genomic alignments
Gene tree
Gene gain/loss tree
Orthologues
Paralogues
Ontologies
GO: Anatomical entity
Phenotypes
Genetic Variation
Variant table
Variant image
Structural variants
Gene expression
Pathway
Molecular interactions
Regulation
External references
Supporting evidence
ID History
Gene history
Configure this page
Custom tracks
Export data
Share this page
Bookmark this page
Gene: ABCA12 ENSG00000144452
Description
ATP binding cassette subfamily A member 12 [Source:HGNC Symbol;Acc:HGNC:14637]
Gene Synonyms
DKFZP434G232, ICR2B, LI2
Location
Chromosome 2: 214,931,542-215,138,626 reverse strand.
GRCh38:CM000664.2
About this gene
This gene has 3 transcripts (splice variants), 216 orthologues, 11 paralogues and is associated with 6 phenotypes.
Transcripts
Show transcript table
Summary
Name
ABCA12 (HGNC Symbol)
MANE
This gene contains MANE Select ENST00000272895, ENSP00000272895
UniProtKB
This gene has proteins that correspond to the following UniProtKB identifiers: Q86UK0
RefSeq
This Ensembl/Gencode gene contains transcript(s) for which we have selected identical RefSeq transcript(s). If there are other RefSeq transcripts available they will be in the External references table
CCDS
This gene is a member of the Human CCDS set: CCDS33372.1, CCDS33373.1
Ensembl version
ENSG00000144452.15
Other assemblies
There is no ungapped mapping of this gene onto the GRCh37 assembly.
View this locus in the GRCh37 archive: ENSG00000144452
Gene type
Protein coding
Annotation method
Annotation for this gene includes both automatic annotation from Ensembl and Havana manual curation, see article.
Go to Region in Detail for more tracks and navigation options (e.g. zooming)
Add/remove tracks
Custom tracks
Share
Resize image
Export image
Reset configuration
Reset track order
227.09 kb
Forward strand
214.94Mb 214.96Mb 214.98Mb 215.00Mb 215.02Mb 215.04Mb 215.06Mb 215.08Mb 215.10Mb 215.12Mb 215.14Mb
Genes (Primary set from GENCODE 48)
SNHG31-205 - ENST00000670391 >
lncRNA
ENST00000628464 >
lncRNA
Contigs
AC114780.3 >
AC072062.3 >
AC072062.3 >
Genes (Primary set from GENCODE 48)
< ABCA12-201 - ENST00000272895
protein coding

### Gene ABCA4 Details

It offers a comprehensive overview of all clinically relevant variants linked to a specified gene. It provides detailed metrics for each variant, including chromosomal position, HGVS c. and p. notation, allele frequencies, carrier rates, and estimated metrics for at-risk couples. By combining gene-level statistics with specific variant-level data, this page enables an in-depth analysis of how particular genes contribute to disease, thus supporting both clinical interpretation and research efforts.

**Gene:** ABCA4

**Disorder:** Retinal dystrophy

**Allele Frequency:** 0.0221943

**Carrier Rate:** 0.0434034

**Max At-Risk Couples rate:** 0.00188386

The table below lists all clinically relevant variants identified in the **ABCA4** gene based on the Emirati cohort dataset.

| id | Chromosome Position | Gene Name | HGVS c. (Clinically Relevant) | HGVS p. (Clinically Relevant) | Allele Count | Allele Frequency |
|---|---|---|---|---|---|---|
| 3 | 94473807 | ABCA4 | NM_000350.3:c.5882G>A | NP_000341.2:p.Gly1961Glu | 40 | 0.01675% |
| 41 | 94520684 | ABCA4 | NM_000350.3:c.2570T>C | p.Leu857Pro | 7 | 0.002931% |
| 162 | 94510248 | ABCA4 | NM_000350.3:c.2971G>C | NP_000341.2:p.Gly991Arg | 2 | 0.000838% |
| 349 | 94490577 | ABCA4 | NM_000350.3:c.4567C>T | p.Gln1523Ter | 1 | 0.000419% |
| 350 | 94508353 | ABCA4 | NM_000350.3:c.3292C>T | NP_000341.2:p.Arg1098Cys | 1 | 0.000419% |
| 351 | 94526295 | ABCA4 | NM_000350.3:c.1958G>A | NP_000341.2:p.Arg653His | 1 | 0.000419% |
| 352 | 94578618 | ABCA4 | NM_000350.3:c.71G>A | NP_000341.2:p.Arg24His | 1 | 0.000419% |

2. Downloadable Data Tables (AVDB Data Exports)
   These curated datasets are derived from clinical exome analysis of 1,194 unrelated Emirati individuals and can be freely downloaded from the AVDB portal.
   - S1. Comprehensive variant allele frequency table derived from diagnostic exome data of 1,194 Emirati individuals: https://avdb-arabgenome.ae/downloads/download.php?id=4&type=xlsx
   - S2. Summarized carrier frequencies per gene across the AVDB cohort for disease risk estimation: https://avdb-arabgenome.ae/downloads/download.php?id=3&type=xlsx
   - S3. List of genes included in the UAE's official premarital carrier screening program: https://avdb-arabgenome.ae/downloads/download.php?id=2&type=csv
   - S4. Full list of disorders represented in AVDB, mapped to their associated genes and inheritance patterns: https://avdb-arabgenome.ae/downloads/download.php?id=5&type=xlsx
